# Integrating regulatory DNA sequence and gene expression to predict genome-wide chromatin accessibility across cellular contexts

**DOI:** 10.1101/605717

**Authors:** Surag Nair, Daniel S. Kim, Jacob Perricone, Anshul Kundaje

## Abstract

**Motivation:** Genome-wide profiles of chromatin accessibility and gene expression in diverse cellular contexts are critical to decipher the dynamics of transcriptional regulation. Recently, convolutional neural networks (CNNs) have been used to learn predictive cis-regulatory DNA sequence models of context-specific chromatin accessibility landscapes. However, these context-specific regulatory sequence models cannot generalize predictions across cell types.

**Results:** We introduce multi-modal, residual neural network architectures that integrate cis-regulatory sequence and context-specific expression of trans-regulators to predict genome-wide chromatin accessibility profiles across cellular contexts. We show that the average accessibility of a genomic region across training contexts can be a surprisingly powerful predictor. We leverage this feature and employ novel strategies for training models to enhance genome-wide prediction of shared and context-specific chromatin accessible sites across cell types. We interpret the models to reveal insights into cis and trans regulation of chromatin dynamics across 123 diverse cellular contexts.

**Availability:** The code is available at https://github.com/kundajelab/ChromDragoNN

## 1 Introduction

Cost-effective, sequencing-based functional genomics assays such as RNA-seq, ChIP-seq, DNase-seq and ATAC-seq have enabled large-scale profiling of epigenomes and transcriptomes across diverse cellular contexts (Consortium, 2012; Kundaje *et al*., 2015). These datasets provide a unique resource to understand the relationship between regulatory DNA sequence, chromatin state and gene expression.

DNase-seq (Thurman *et al*., 2012; Boyle *et al*., 2008) or ATAC-seq (Buenrostro *et al*., 2013) experiments profile the accessible chromatin landscape typically bound by regulatory DNA binding proteins such as transcription factors (TFs). Chromatin accessibility is highly dynamic across cellular contexts (Thurman *et al*., 2012). Chromatin accessibility of a regulatory element is largely a function of the combinatorial cis-regulatory code of TF binding sequence motifs embedded in its DNA as well as the availability and activity of the trans-regulatory proteins such as TFs that bind them.

A large body of literature has focused on developing computational models to decipher the cis-regulatory sequence code of cell-type specific chromatin accessibility landscapes. Recently, convolutional neural networks (CNNs) have been used to learn the cis-regulatory grammars encoded in regulatory DNA sequences associated with cell-type specific *in vivo* TF binding and chromatin accessibility (Kelley *et al*., 2016; Alipanahi *et al*., 2015a; Quang and Xie, 2016; Haoyang Zeng and Gifford, 2016; Zhou and Troyanskaya, 2015; Alipanahi *et al*., 2015b). By learning a series of de-novo motif-like pattern detectors (called convolutional filters) and non-linear activation transformations, CNNs are able to map raw DNA sequence across the genome to binary or continuous measures of associated regulatory activity profiles without explicit feature engineering.

The Basset model (Kelley *et al*., 2016) is a state-of-the-art CNN architecture that predicts binary chromatin accessibility in a specific cellular context across the genome as a function of local 600 bp DNA sequence context around each bin. The Basset model is also a multi-task architecture trained simultaneously on binary chromatin accessibility profiles from multiple cellular contexts (each context is a prediction task) and produces a vector of outputs for any genomic position containing the probability of accessible chromatin state at that position in each of the cellular contexts (task). The input DNA sequences represented using a one-hot encoding is transformed by three convolution layers. A rectified linear unit (ReLU) nonlinear transformation is applied to the output of the final convolution layer and a pooling operation takes the maximum across a window of adjacent positions. These transformations are then passed to three fully connected layers followed by a logistic non-linearity for each task (cellular context) that outputs the probability of accessibility. The convolutional filters learned by Basset were visualized and interpreted to infer putative cis-regulatory sequence drivers of context-specific chromatin accessibility. The model was also used to score putative regulatory genetic variants using an in-silico mutagenesis approach.

The Basset model was recently enhanced by factorizing the convolution layers (Wnuk *et al*., 2017) (Factorized model). The Factorized model increases the model depth - the 3 convolution layers in Basset are replaced by 9 convolution layers. Further, the first two convolution layers in Basset which contain convolutional filters (motif-like pattern detectors) of widths 19 and 11 respectively are factorized into multiple convolution layers with smaller widths. The authors note that these modifications enhance prediction performance and reduce learning time.

While these and other sequence-only models (Zhou and Troyanskaya, 2015; Kelley *et al*., 2018) have provided useful insight into context-specific cis-regulatory sequence features and the context-specific impact of regulatory genetic variants, these models cannot be used to predict chromatin accessibility or other regulatory profiles in cellular contexts not present in the training set. This is largely because these sequence-only models do not model the regulatory activity of trans-factors that vary across cellular contexts. Gene expression levels of trans-factors as measured by RNA-seq provide a useful, albeit indirect surrogate for their availability and activity in different cellular contexts. Models that can integrate cis-regulatory DNA sequence and trans-regulator expression should in principle be able to generalize to predict chromatin accessibility landscapes across cellular contexts. Such a model would be very valuable because it would enable prediction of chromatin accessibility profiles in large collections of cellular contexts that are currently characterized only by RNA-seq (Collado-Torres *et al*., 2017). Moreover, interpreting such an integrative model would also provide insights into cis-regulatory sequence features and trans regulators that are predictive of chromatin dynamics across cellular contexts.

Deep-learning architectures allow this kind of flexibility to integrate multi-modal data i.e. DNA sequence coupled with RNA expression profiles. Hence, we expand upon previous work to predict genome-wide maps of chromatin accessibility using sequence and gene expression data (Kelley *et al*., 2016; Wnuk *et al*., 2017). We introduce multi-modal, residual neural network architectures (He *et al*., 2016) that integrate cis-regulatory sequence and context-specific expression of trans-regulators to predict genome-wide chromatin accessibility profiles across cellular contexts. We show that the average accessibility of a genomic region across training contexts can be a powerful baseline predictor. We leverage this feature and employ novel strategies for training models to enhance prediction performance of shared and context-specific chromatin accessible sites across cell types. Further, we show that we can interpret these cross-cell type models to reveal insights into cis and trans regulators of chromatin dynamics across 123 diverse cellular contexts.

## 2 Methods

### 2.1 Chromatin accessibility data

DNase-seq datasets profiling genome-wide chromatin accessibility were downloaded from the Roadmap Epigenomics Project^1^ and ENCODE^2^. The complete list of DNase-seq datasets and their identifiers is provided in Supp. Table 1. The fastq files were aligned with BWA aln (v0.7.10), where all datasets were treated as single-end, with ENCODE default alignment parameters. After mapping, reads were filtered to remove unmapped reads and mates, non-primary alignments, reads failing platform/vendor quality checks, and PCR/optical duplicates (-F 1804). Low quality reads (MAPQ *<* 30) were also removed. Duplicates were then marked with Picard MarkDuplicates and removed. The final filtered file was then converted to tagAlign format (BED 3+3) using bedtools bamtobed. Cross-correlation scores were then obtained for each file using phantompeakqualtools (v1.1).

All files were checked to have cross-correlation with a quality tag above 0 and discarded if not. For the ENCODE data generated from the Stam Lab protocol, all datasets were trimmed to 36 bp and then combined if technical replicates. Read depths were considered, and a standardized depth of 50 million reads was set for the final datasets. As such, the files were filtered to remove mitochondrial reads, filtered for mappability, and then subsampled to 50 million reads. For the ENCODE data generated from the Crawford Lab protocol, the same procedure as above was performed, except reads were trimmed to 20 bp due to the different library generation protocol. For the Roadmap data, which was all generated by the Stam Lab protocol, the same procedure as above was performed with trimming to 36 bp, and files were only combined to give a minimum read depth of 50 million reads, since each file came from a different developmental time point. These trimmed, filtered, subsampled tagAlign files were then used to generate signal tracks and call peaks. Signal tracks and peaks were called with a loose threshold (*p <* 0.01) with MACS2 to generate bigwig files (fold enrichment and *p*-value) and narrowPeak files, respectively.

For reproducible peak sets, we performed pseudoreplicate subsampling on the pooled reads across all replicates (taking all reads from the final tagAligns and splitting in half by random assignment to two replicates) and retaining reproducible peaks passing an Irreproducible Discovery Rate (IDR v2.0.3) (Li *et al*., 2011) (https://github.com/kundajelab/idr) threshold of 0.1 to get a reproducible peaks for each DNase experiment. The pipeline is available in a Zenodo record https://doi.org/10.5281/zenodo.156534.

We bin the human genome (GRCh37 assembly) into 200 bp bins (*i*) every 50 bp. For each of the 123 cellular contexts (*j* = {1 … 123}), all bins are assigned binary labels (*y*_*i,j*_ ∈ {0, 1}) corresponding to accessible (+1) or inaccessible (0) state based on whether they overlap (> 50% overlap) context-specific reproducible DNase-seq peaks or not. The genome-wide binary labels for each task *j* (cellular context) are highly imbalanced (Proportion of positive bins: min=3%, median=7%, max=10% across cell types). The complete binary label matrix is available via a Zenodo archive https://doi.org/10.5281/zenodo.2603199. The cis-regulatory sequence context (*S*_*i*_) for each bin *i* is represented using 1000 bp of genomic DNA sequence centered at the bin. We use a 1000 bp sequence context since previous work showed performance gains using contexts up to 1000bp (Zhou and Troyanskaya, 2015; Avsec *et al*., 2018).

### 2.2 Gene expression data

RNA-seq fastq files (no subsampling, no filtering, no trimming) from Roadmap and ENCODE were mapped using the STAR aligner (version 2.4.1d), using ENCODE default parameters. GENCODE release 19 (GRCh37.p13) transcriptome annotations were used. To determine the strandedness of the file (which is needed for RSEM quantification), the infer_experiment.py script from RSeQC (version 2.6.4) was used in conjunction with the STAR output that was sorted by coordinate. The strandedness and the pairedness (paired end or single end) of the experiment were passed on to RSEM (version 1.2.21). For RSEM, we used ‘-estimate-rspd’ to estimate read start position distribution, and we did not calculate confidence bounds. If the experiment was stranded, we set ‘–forward-prob’ to be 0, and unstranded experiments were left at default. The transcriptome aligned file from STAR was used in the RSEM run. The complete list of RNA-seq datasets and their identifiers is provided in Supp. Table 3. The pipeline is available at https://github.com/ENCODE-DCC/rna-seq-pipeline (v1.0).

The final dataset includes RNA-seq data associated with each of the 123 cell types. We extract the TPM (transcripts per million) values and normalize the values by taking the log of the values.

The trans-regulatory feature space *R*_*j*_ for each cellular context *j* = {1 … 123} is represented by the log(TPM) expression levels of a list of 1630 putative TFs as curated by the FANTOM5 consortium ^3^ of human TFs. The TF gene expression feature matrix is available via a Zenodo archive https://doi.org/10.5281/zenodo.2603199.

### 2.3 ChromDragoNN Neural network architecture

Our goal is to learn a model *F*(*S*_*i*_, *R*_*j*_) that can predict the binary chromatin accessibility state *y*_*i,j*_ at any bin *i* in genome in any cellular context *j* as a function of the one-hot encoded 1 Kb cis-regulatory sequence context *S*_*i*_ of bin *i* and the expression of 1630 TFs *R*_*j*_ in cellular context *j*. We use a multi-modal neural network model to integrate the cis-sequence and trans-expression modalities and optionally the mean accessibility of the bin across cell types.

The one-hot encoded sequence *S*_*i*_ for each bin *i* in the genome is fed into a residual convolutional neural network (ResNet) model (Fig. 1A). The ResNet architecture includes hierarchically arranged convolution layers that are able to map one-hot encodings of raw DNA sequence input data to learn complex representations without explicit feature engineering. Each convolution layer learns and scans a set of weight matrix pattern detectors (convolutional filters) across its input and detects patterns in the input sequence. Residual neural networks (ResNets) (He *et al*., 2016) have been show to be more effective for training CNNs with a large number of layers by introducing skip connections between blocks of convolution layers to optimize gradient flow and improve learning. Utilizing these concepts, we use a ResNet architecture to extend previous models (Kelley *et al*., 2016; Wnuk *et al*., 2017). The residual network (He *et al*., 2016) consists of blocks in which the input is transformed through one or more convolutional layers to an intermediate output to which the input is added back. In our model, the convolution layers within a block preserve the input dimensions.

**Fig. 1:**
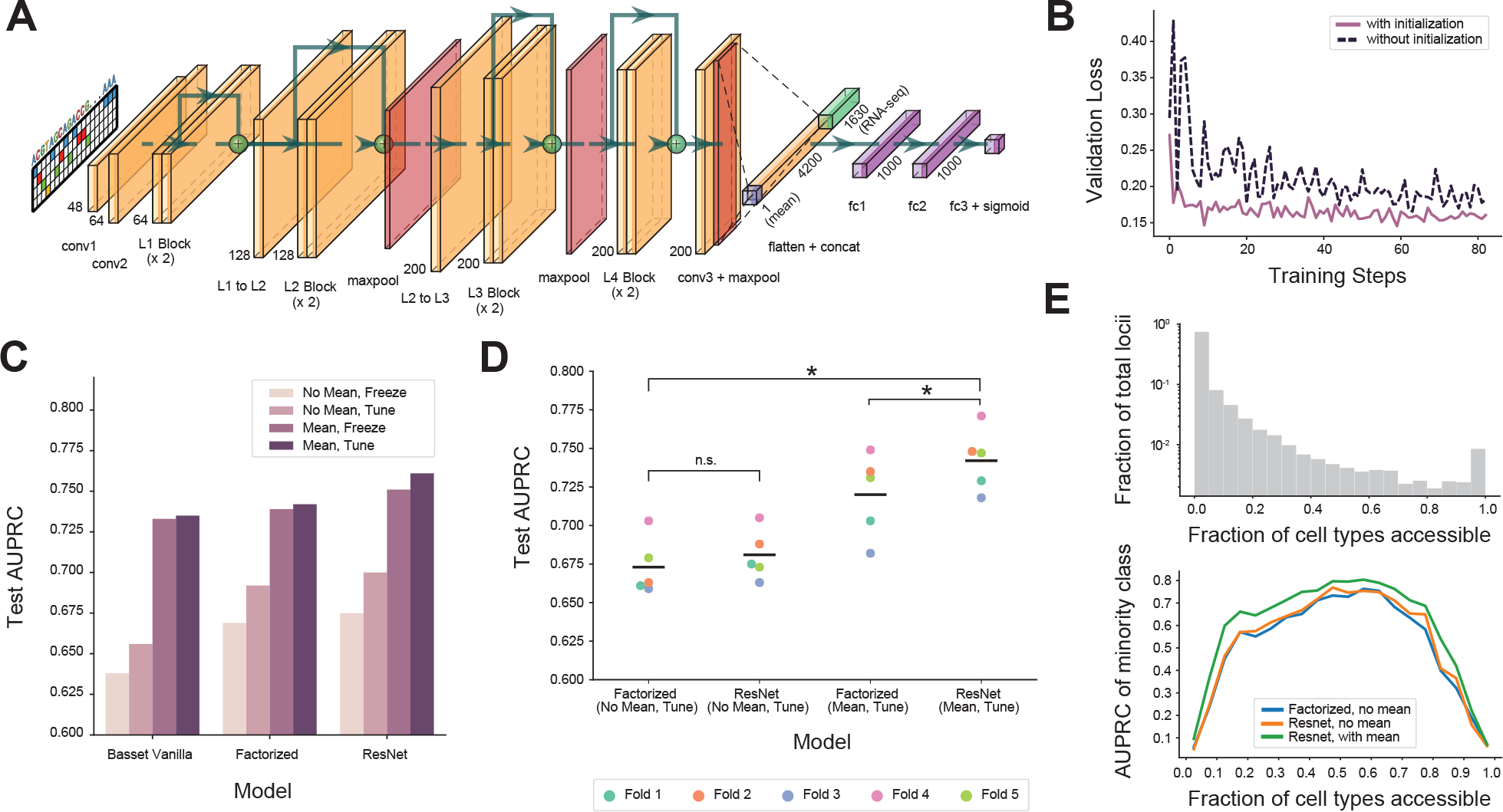
Improved training methods and new architecture design enhances model performance. **(a)** Model architecture for the ResNet model. The RNA-seq inputs and mean accessibility (if used) are concatenated after the convolutional layers. **(b)** The validation set loss over training steps for a model (Basset architecture for sequence mode) with and without two-stage learning (without mean accessibility as an input feature). In two-stage learning the weights of the convolutional layer of the model are initialized from a model first trained to map sequence to chromatin accessibility for all training cell types. **(c)** The test set AUPRC of the original Basset model, Factorized model, and ResNet model under 4 training paradigms: with and without mean accessibility as an input feature, and with (Tune) and without (Freeze) fine-tuning convolution layers in 2^nd^ stage. Numbers reported on a fixed training, validation and test split with 103 training, 10 validation and 10 test cell types. Models using mean accessibility as an input feature significantly outperform models without mean accessibility. **(d)** 5-fold cross validation performance of the ResNet model compared to the Factorized model with and without mean locus accessibility as an input to the model. Each fold contains a split over 123 cell types in the dataset. All models trained using 2-stage scheme with all weights tunable in second stage. Wilcoxon signed rank test (single-tailed) was performed with *n* = 5, n.s. not significant, \**P* < 0.05. **(e)** Binned AUPRC of Factorized model without mean accessibility, ResNet model without mean accessibility, and ResNet model with mean accessibility. Loci are binned by the fraction of training cell types that are accessible, and AUPRC is computed for predictions on test cell types for each bin. Note that AUPRC is computed for the minority class-when fraction of accessible cell types *>* 0.5, AUPRC is computed on non-accessible regions. Gray bars indicate the fraction of loci having a certain fraction of accessible cell types. Numbers reported on a training, validation and test split same as for (c).

To provide the model with quantitative information on the availability of trans-regulator TFs, we follow recent work (Wnuk *et al*., 2017) that extended the Basset model to predict chromatin accessibility in held-out cellular contexts, using RNA-seq profiles as surrogates of cell-type specific availability and activity of trans-regulators. RNA-seq profiles have been shown to uniquely identify individual cell types while preserving biological similarity between cell types (Sudmant *et al*., 2015). We use log(TPM) RNA expression levels of 1630 transcription factors as a meaningful representation of trans-regulatory cell state, as TFs are the DNA binding proteins that would affect chromatin accessibility by binding cis-regulatory sequence patterns. The sequence ResNet-CNN component of the model learns cis-regulatory sequence patterns and returns a transformed sequence-based feature space as intermediate representation. The TF RNA-seq vector *R*_*j*_ for cellular context *j* is concatenated with this intermediate sequence representation, which is then passed through fully connected neural network layers and a logistic non-linearity to produce an output *F*(*S*_*i*_, *R*_*j*_) representing the predicted probability that the bin *i* is accessible in the cellular context *j*. The mean accessibility for bin *i* across all training cell types, if used, is concatenated at the final fully connected layer. The complete sequential network is as follows:

One-hot input sequence of dimension 1000. 2 convolutional layers with 48 and 64 channels respectively, filter size (3,1). 2 residual blocks, each with 2 convolution layers with 64 channels and filter size (3,1). 2 residual blocks, each with 2 convolution layers with 128 channels and filter size (7,1). 2 residual blocks, each with 3 convolution layers with 200 channels and filter sizes (7,1), (3,1), (3,1) respectively. 2 residual blocks, each with 2 convolution layers with 200 channels and filter size (7,1). The output is flattened and concatenated with gene expression. In case of mean accessibility models, the mean is concatenated. Fully connected layer with 1000 dimension output. Fully connected layer with 1000 dimension output. Fully connected layer with 1 output dimension.

A single convolution layer is present after each residual block (except the third) to transform the number of channels. Batch normalization (Ioffe and Szegedy, 2015) layers are present after each layer. A max pool is applied after the last 3 residual blocks. We use the ReLU non-linearity transform. We use a fixed dropout of 0.3 on the fully connected layers.

### 2.4 Multi-stage training

We randomly split our 123 cellular contexts into training, validation and test sets across 5 folds (Supp. Table 2). For each fold, we train models genome-wide across the training cell types. The validation set cell types are used for hyperparameter tuning. The models are evaluated based on their genome-wide predictions in the held-out cell types in the test sets.

The shift from a multi-task cell type specific sequence-only model to a single-task, cross-cell type, multi-modal model increases the number of training examples by a factor of *C*, equal to the number of cell types in the training data. The increased size of the training data has implications for training. A naive training setup could potentially take up to a factor *C* longer to train compared to a fixed cell type model. To improve efficiency, performance and interpretability, we train our models in two steps: the first stage pre-trains a multi-task sequence-only model that maps sequence of each genomic bin to accessibility labels in each of the cellular contexts in the training set as individual tasks. The second stage trains the multi-modal model across all genomic bins and cellular contexts in the training set by initializing the sequence-mode’s convolutional layer weights using the pre-trained model. The two-stage training scheme provides added flexibility in that during the second stage of training, the convolutional layer weights may or may not be frozen while the fully connected layers are trained.

### 2.5 Model Training and testing

We use the Adam optimizer (Kingma and Ba, 2014) on binary cross entropy loss to update our network’s weights, along with batch normalization on the convolution and fully connected layers. We use the default PyTorch v0.4 parameter initialization method (LeCun *et al*., 2012). We perform hyperparameter searches for all stage 1 models with batch sizes (128, 256) and learning rates (2e-2, 2e-3, 2e-4), and for stage 2 models with batch sizes (256, 512, 1024) and learning rates (1e-3, 1e-4). To mitigate the class imbalance, we maintained a 1:3 ratio of positives to negatives per batch by upsampling accessible regions in the second stage of training.

Given the significant class imbalance in the labels, we use the area under precision-recall curve (AUPRC) as our primary performance evaluation measure.

### 2.6 Motif extraction

The dynamics of chromatin accessibility of regulatory elements across cellular contexts is a result of distinct subsets of context-specific TFs binding combinations of motifs encoded in the sequence of the regulatory elements (Sherwood *et al*., 2014; Voss and Hager, 2014). Deep neural network models of regulatory DNA sequence implicitly learn these motifs as distributed representations across the convolutional filters. Hence, valuable insights on predictive regulatory sequence features can be obtained by interpreting the model. A commonly used approach for involves directly visualizing the convolutional filters or deriving position weight matrices from subsequences that maximally activate filters (Kelley *et al*., 2016). However, this approach has the drawback that the motifs obtained from individual filters are often redundant or incomplete since the models learn distributed representations (Shrikumar *et al*., 2018). An alternative approach is to use feature attribution methods to interpret predictive patterns in specific input DNA sequences. These feature attribution methods (Shrikumar *et al*., 2017; Sundararajan *et al*., 2017; Simonyan *et al*., 2013) decompose the output prediction of a model for a specific input sequence of interest in the form of contribution scores of individual nucleotides in the sequence. Nucleotides with high positive scores can be interpreted as driving the prediction for the sequence. Feature attribution methods allow for instance-by-instance interpretation of predictive patterns but do not provide a global summary of predictive motifs across all accessible sites within and across cellular contexts. Hence, we used a new method we recently developed called TF-MoDISco (v0.2.1) (Shrikumar *et al*., 2018) that (i) identifies predictive sequence patterns within the sequences of each accessible site across the genome in a cell context of interest as subsequences (called seqlets) with significant contribution scores derived using a feature attribution method (specified below); (ii) computes a similarity matrix between all predictive seqlets across the accessible landscape; and (iii) clusters the seqlets into non-redundant motifs. To obtain nucleotide-resolution contribution scores for each input sequence corresponding to accessible bins in the genome in a specific cellular context, we used the gradient of the logit of the output probability of the model (predicted probability of site being accessible in the specific cellular context) with respect to the one-hot DNA sequence, gated by the observed nucleotides in the input sequence. To focus on motifs associated with dynamic chromatin accessible sites, for each cellular context, we extracted the contribution score profiles from the ResNet model (that does not use mean accessibility as an input feature) for subsets of 20,000 bins that are accessible in the given cellular context and in < 30% of all the cell types. Contribution score profiles computed for these 20K sequences in each cellular context were passed to TF-MoDISco to learn context-specific globally predictive motifs. The TF-MoDISco motifs were matched against a database of known TF motifs using Tomtom (Gupta *et al*., 2007).

## 3 Results

### 3.1 Accurate prediction of chromatin accessibility across cellular contexts from DNA sequence and gene expression with multi-stage training

We developed multi-modal neural network architectures to predict the binary chromatin accessibility state at each bin in the genome in any cellular context by integrating 1Kb cis-regulatory sequence context around each genomic bin and gene expression levels of 1630 transcription factors in the specific cellular context. Models were trained on a subset of training cell types and their performance was reported based on genome-wide predictions in held-out test cell types. We developed a two-stage learning strategy to improve efficiency, performance and interpretability of the models. In the first stage, we pretrained a multi-task sequence-only model across all training cell types. In the second stage, we trained a multi-modal model integrating sequence and expression, where we initialized the convolutional layer weights of the sequence model from the first stage. We found that tuning the convolution layers in the second stage consistently improved performance over freezing the weights of the layers at an increased computational cost. Further, pre-training the sequence mode consistently improved training time and performance (Fig. 1B).

We experimented with different CNN architectures, training strategies and tested the impact of adding an additional feature - the mean accessibility of a genomic bin across training cell types. After evaluating the various models on our dataset, our best model achieves an AUPRC = 0.76 and AUROC = 0.954, outperforming previously published model architectures trained and tested on matching data (AUPRC =0.69, AUROC = 0.937) (Fig. 1C).

### 3.2 Using mean accessibility as an input feature boosts performance

A key difference between cell-type specific models and cross-cell type models is that cross-cell type models can make use of statistics based on the accessibility state of each genomic bin (locus) across the training cell types. For each bin in the genome, we computed the mean of the binary accessibility values across all cell types in the training set. Since binary accessibility is 0 if the locus is closed and 1 if open, the mean accessibility is a value in [0, 1] that is equivalent to the fraction of cell types in which the bin is accessible.

We observed that mean accessibility is a strong baseline predictor of chromatin accessibility across cell types (also recently reported by Schreiber *et al*. (Schreiber *et al*., 2019)). Setting the predicted accessibility of a locus equal to its mean accessibility across training cell types yielded an AUPRC of 0.579 and an AUC of 0.902 on the test set. This method is oblivious to the test cell type and in fact assigns the same values to all test cell types for a given bin. A stronger baseline is achieved by computing a weighted average of accessibility across training cell types, where the weight is proportional to the similarity between RNA-seq profiles of the training and test cell types. The resulting predictions yield an AUPRC of 0.587 and AUC of 0.903 which are marginally better than the unweighted version.

All our multi-modal models that use sequence and expression substantially outperform this strong baseline predictor (mean-baseline AUPRC=0.579, weighted-mean baseline AUPRC=0.587, Basset+expr AUPRC=0.656, FactorizedBasset+expr AUPRC=0.692, ResNet+expr AUPRC=0.700). However, we decided to capitalize on the strong mean baseline and decided to use it as an auxiliary input feature to the multi-modal model. The single scalar mean accessibility value for each bin is concatenated with the output of first feed forward layer. We observe substantial improvements when the mean accessibility feature is provided as an input to the model (Fig. 1 C,D). Across 3 different types of architectures that we trained, incorporating the mean as an input feature improves the performance of the model by as much as 0.09 AUPRC.

### 3.3 Residual network architecture outperforms previous architectures

Residual neural networks (ResNets) (He *et al*., 2016) have been shown to be highly effective for training deeper CNNs with a large number of layers. ResNets provide added flexibility to CNNs by introducing skip connections between blocks of convolution layers. In practice, while the performance of ordinary CNNs saturates or even drops with increasing layers (Srivastava *et al*., 2015), ResNets have made possible training of CNNs often having more than 100 convolution layers. ResNets have also recently been used to train high performance deep learning sequence models of splicing (Jaganathan *et al*., 2019).

We implemented a ResNet architecture that uses 23 convolution layers across 8 residual blocks. Following the Factorized model, we used convolution filters with shorter widths. (Fig. 1D) shows the results of a 5-fold cross validation performed on our dataset. We compared the performance of the model with the Factorized Model with and without passing mean locus accessibility as an input to the model. In both cases, the ResNet architecture improved upon the performance of the Factorized model. Overall, our best performing ResNet(+mean accessibility) model achieves an AUPRC of 0.76 while the the previous best published model in the literature i.e. the Factorized Basset model (Wnuk *et al*., 2017) achieves 0.69 (Fig. 1C) on a matched training/validation/test data split.

Next, in order to understand performance variation as a function of cell type specificity of accessible sites, we grouped genomic bins based on the fraction of cell types in which bins exhibit accessibility. For each group, we compared the AUPRC of our best ResNet model that included mean accessibility as auxiliary input with the previous best published model i.e. Factorized Basset without mean accessibility (Fig. 1E). Our models consistently outperform the previous state-of-the-art across all groups.

### 3.4 Model interpretation reveals cell type specific cis-regulatory sequence features and associated trans-regulators

Understanding what the model is utilizing in the DNA sequence input is of interest, and previous work has successfully shown that CNNs learn predictive motif-like patterns of cell-type relevant TFs from regulatory DNA sequences (Kelley *et al*., 2016). However, the model learns a distributed representation of the sequence features. Hence, interpreting individual convolutional filters results in redundant and partially complete motifs. Instead, we use TF-MoDISco, a new method we recently developed, for distilling consolidated motifs from sequence-based deep learning models (Shrikumar *et al*., 2018). First, we use a feature attribution approach (gradient *×* input) to infer contribution scores attributed by the model to each nucleotide in chromatin accessible sequences in each cellular context, to the output prediction associated with the sequence. Predictive nucleotides and motif instances get highlighted with high positive contribution scores. The same sequence can have different contribution score profiles across different cellular contexts representing dynamic regulation of the region by different sequence motifs (Fig. 2A). For each cellular context, we sample a subset of bins that are labeled accessible, obtain contribution scores for corresponding input sequences and extract motifs using TF-MoDISco with default parameters. The motifs are matched against a database of known motifs of TFs using Tomtom (Gupta *et al*., 2007). The sets of motifs retrieved for each cellular context reflect the globally predictive TF motif patterns learned by the model for that context (Fig. 2B).

**Fig. 2:**
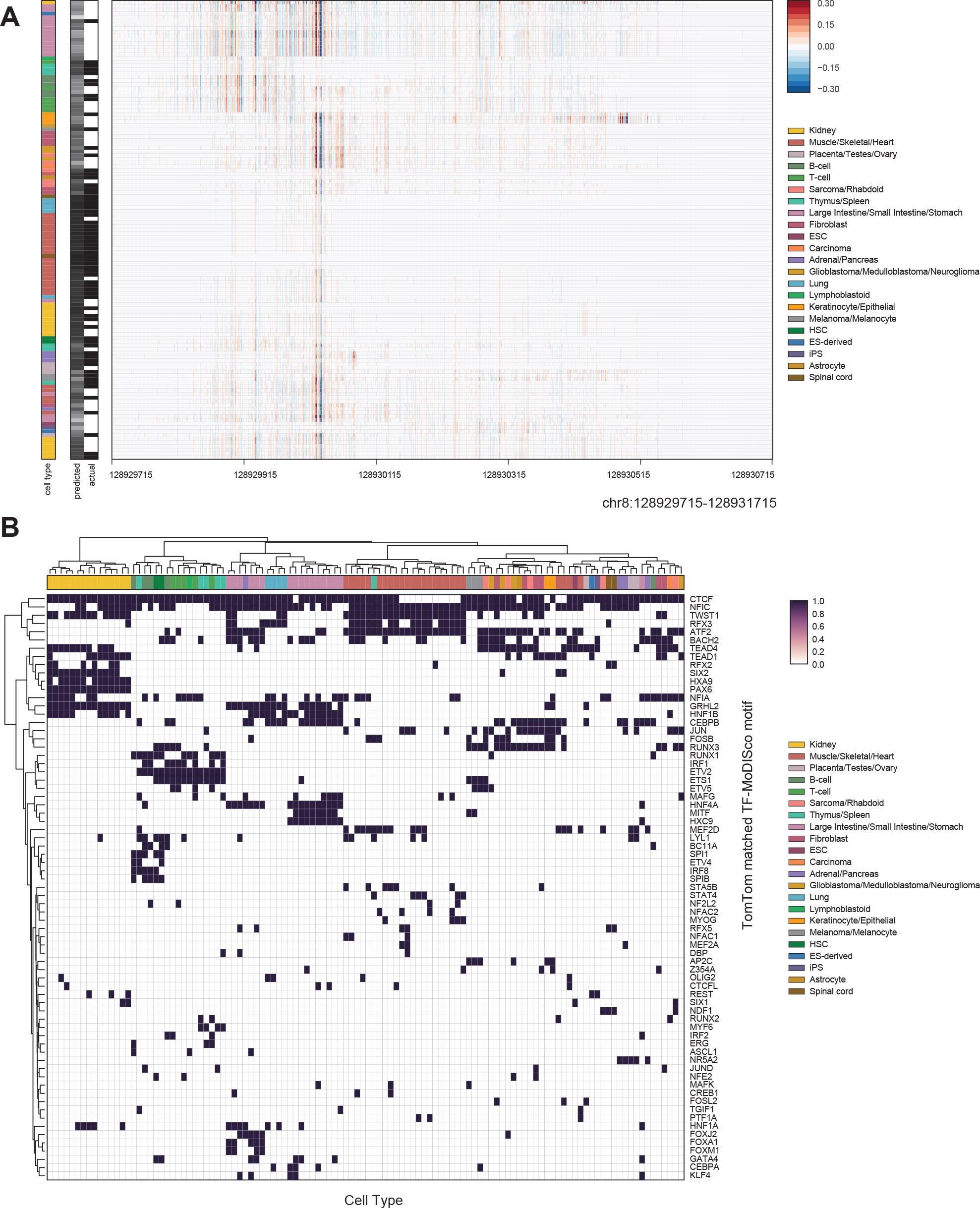
Cell-type specific transcription factor motifs distilled from the ResNet model. **(a)** Gradient × input contribution scores of each nucleotide (columns) in an example genomic sequence (chr8:128929715-128931715) across different cellular contexts (tasks shown as rows). The obtained nucleotide resolution contribution scores for the same genomic sequence can differ between cell-types reflecting differential chromatin accessibility and differences in regulation of the sequence, as shown in this example locus. **(b)** Summary of motifs learned by the model for individual cell types. The TF-MoDISco method is used to distill consolidated motifs learned by the model for each cell type using a subset of sequences which are accessible in the respective cell type. The returned motifs are then matched to known motifs of TFs using Tomtom (Gupta *et al*., 2007).

The model learned known DNA motifs of ubiquitous as well as cell-type specific TFs that match the canonical roles of TFs in different lineages (Fig. 2B). As reported in (Kelley *et al*., 2016), the model learns the CTCF motif as a widely important sequence element for accessible regions across cellular contexts (Ong and Corces, 2014). The HNF1A and HNF4A motifs are more narrowly predictive of accessibility in hepatocyte-related, large and small intestinal contexts (D’Angelo *et al*., 2010). The model discovers SIX2 motif as a key predictor in kidney-related contexts (Kobayashi *et al*., 2008). TWIST1 motif is retrieved for contexts of mesenchymal origin (Qin *et al*., 2012), while RUNX1, ETS1 and IRF1 motifs are mainly discovered only in specific hematopoietic cell types (Brien *et al*., 2011). GRHL2 motif is discovered in the lung, epithelial cells, and kidneys, which matches known differential expression patterns of GRHL family transcription factors across cell types (Aue *et al*., 2015). No prior information about sequence motifs is provided to the model, suggesting that the model is effective at extracting cell context relevant cis-regulatory features from the DNA sequence input.

Many of the discovered motifs are cell-type specific, which suggested that intersecting these results with the dynamics of RNA expression profiles of trans-regulators could potentially lead us to the TFs that potentially bind these discovered motifs. For each discovered motif, we determined all the TFs (often from the same family) that could potentially bind the motif. We computed the binary vector of dynamic motif activity for each motif across cell types (whether that motif was discovered by TF-MoDISco in the cell type or not). We computed the Pearson correlation between the motif activity vector and the vector of expression levels of matching TFs across those cell types. We show the top 15 most correlated TFs in Fig. 3A. This analysis highlighted several known key regulators, both universal and cell-type specific, across a variety of cell types. TWIST1 is a known regulator in mesenchymal cell types and is highlighted as important in muscle cell types and fibroblasts (Qin *et al*., 2012). RUNX3 and IRF1 are important regulators in blood cell types (Brien *et al*., 2011), while HNF4A is a master regulator in intestinal development (Babeu and Boudreau, 2014). HNF1A, GRHL2, SIX2, and HOXA9 are all regulators known to be important in kidney development (Kobayashi *et al*., 2008; Martovetsky *et al*., 2013; Aue *et al*., 2015), and are highlighted here as important specifically in kidney cell types. Interestingly, ASCL1 is highlighted as important in thymus and spleen cell types, where the expression is also very specifically high in these cell types - this suggests a role for ASCL1 in these cell types that was not elucidated before, though further work is required to fully validate this hypothesis. This analysis thus uncovers possible trans-regulators that modulate cell context-specific chromatin accessibility profiles through predictive cis-regulatory motifs.

**Fig. 3:**
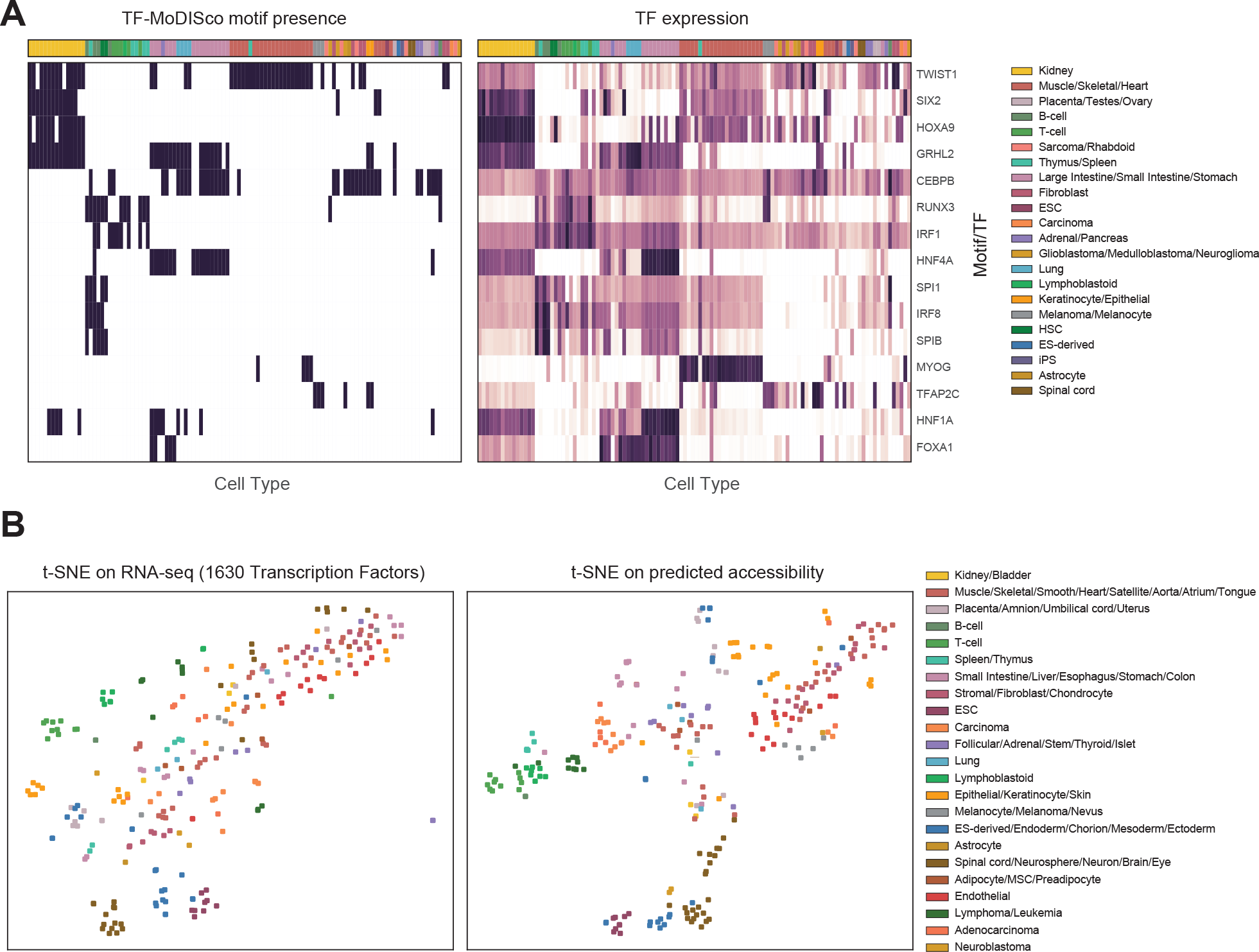
Predictive cis-sequence features and trans-regulators inferred from the models: **(a)** TF-MoDISco motifs (left) and row-normalized log TPM RNA-expression values (right) for each training cell type, where each row is a matching motif and TF. **(b)** t-SNE embedding of 250 additional cell types (points) based on the RNA-seq profiles of 1630 TFs (left) compared to t-SNE embedding of the same cell types based on the predicted chromatin accessibility profiles

### 3.5 Biologically relevant segregation of cell types based on predicted chromatin accessibility

We used our cross-cell type, multi-modal models to impute genome-wide binary chromatin accessibility profiles in 250 additional cellular contexts (See Supp. Table 3) that were not seen in our original dataset and were profiled only using RNA-seq. These new imputed samples were then embedded into 2D visualization using t-SNE (Maaten and Hinton, 2008) to determine how well the imputed accessibility profiles group distinct and related cell types. Comparing an equivalent t-SNE visualization in RNA-seq expression space (using the 1630 TFs as features) to the predicted chromatin accessibility (Fig. 3B), we find that the t-SNE map from imputed accessibility shows improved separation of distinct clusters of samples grouped by cell type and disease state. E.g. the carcinoma cell types and the adenocarcinoma cell types are embedded near each other in the t-SNE from predicted accessibility. Further, the predicted accessibility t-SNE embeds the adenocarcinomas as slightly offset from the carcinomas. While t-SNE embeddings can be unstable and difficult to interpret, our visualizations do suggest that the imputed accessibility profiles do capture biologically meaningful differences and similarities between cell types and that these differences are not simply reflecting differences in expression of the TFs that were used as predictors. This ability to distinguish cell types through imputed accessibility profiles is important because it suggests that given a new expression profile, these models can produce distinct accessibility profiles that may be granular enough to potentially reveal subtypes and finer grained structure beyond the expression profile.

## 4 Discussion

We present an optimized multi-modal residual network architecture that can integrate cis-regulatory DNA sequence and expression of trans-regulators to predict genome-wide binary chromatin accessibility profiles across cellular contexts. The model can be used to predict genome-wide chromatin accessibility in cellular contexts that are only profiled with RNA-seq. This is particularly useful given the large number of profiled transcriptomes that do not have corresponding experimentally profiled epigenomes. We demonstrate that accessibility profiles predicted from sequence and TF expression do not simply recapitulate the landscape of expression profiles across cell types but rather provides a complementary feature space that can discriminate between related and distinct cellular contexts.

Using enhanced training strategies, we achieve a new state of the art in terms of prediction performance across cellular contexts. We show that a two-stage training strategy that pre-trains using only sequence before integrating the expression data improves performance and training time. This method of transfer learning is common in applications in computer vision and natural language processing (Chen *et al*., 2015; Oquab *et al*., 2014). In two-stage model learning, we show that tuning the convolution layers in the second stage offers a benefit over freezing the weights of the layers, however at an increased computational cost. In addition, adding the mean accessibility of a given locus significantly improves performance. Mean accessibility by itself is a surprisingly strong predictor of chromatin accessibility. Combining the mean accessibility with cis-regulatory sequence and trans-regulator RNA expression allows improved prediction performance. Notably, we find that adding mean accessibility as a feature improves performance across all types of accessible sites including the cell type specific and ubiquitously active.

We demonstrate that using a residual CNN architecture for chromatin accessibility prediction results in superior performance compared to previous architectures. Recent related work (Wnuk *et al*., 2017) showed that increasing the number of convolution layers while reducing the width of each convolution layer increases the model performance. Residual neural networks (He *et al*., 2016) allows for connections between non-adjacent layers and have been shown to confer performance gains in deep networks. We observe and confirm similar improvements in model performance for predicting chromatin accessibility models.

Recently developed imputation methods such as ChromImpute(Ernst and Kellis, 2015), PREDICTD (Durham *et al*., 2018) and Avocado (Schreiber *et al*., 2018) also tackle the problem of predicting regulatory profiles in new cellular contexts. However, these frameworks are based on capturing and modeling the local correlation structure between profiles of multiple biochemical markers such as RNA, histone modifications and chromatin accessibility within and across diverse cell types. In our framework, we instead use only one widely available auxiliary modality, the gene expression of trans-regulators. Moreover, the above mentioned imputation methods do not model cis-regulatory DNA sequence and hence lack the ability to interpret biologically meaningful predictive sequence features from the models. Our models enable interpretation of predictive cis-sequence features learned by the models. Using model interpretation methods, we show that our models learn motifs of ubiquitous and lineage specific TFs. Correlating the RNA profiles of TFs with the dynamic predictive activity of motifs discovered by the model provides insights into the TFs that might bind these motifs and the relationship between cis and trans regulatory features.

Our current models predict genome-wide binary chromatin accessibility profiles instead of continuous, quantitative profiles. However, our models can be easily adapted to predict continuous, quantitative profiles at finer resolutions by using regression loss functions Kelley *et al*. (2018). Our models can also be extended to include additional input data modalities or predict other types of genome-wide regulatory profiles such as histone modification profiles. Finally, improved approaches for interpreting multi-modal neural networks will provide significantly more nuanced insights into the complex interactions between cis-regulatory sequence features and trans-regulatory features. More transparent encodings of the gene expression space (e.g. using latent variables that directly model modules of functionally related genes or pathway annotations) would also improve interpretability. Our study highlights the promise of integrative multi-modal deep learning models for learning predictive models that generalize across cellular contexts and obtaining insight into the dynamics of gene regulation.

## Acknowledgments

The computing for this project was performed on the Sherlock and Nautilus clusters. We thank Stanford University, the Stanford Research Computing Center, and the Pacific Research Platform for computational resources.

1 http://www.ncbi.nlm.nih.gov/geo/roadmap/epigenomics/

2 https://www.encodeproject.org/

3 http://fantom.gsc.riken.jp/5/sstar/Browse_Transcription_Factors_hg19

